# Identification of transcription factors regulating senescence in wheat through gene regulatory network modelling

**DOI:** 10.1101/456749

**Authors:** Philippa Borrill, Sophie A. Harrington, James Simmonds, Cristobal Uauy

## Abstract

Senescence is a tightly regulated developmental programme which is coordinated by transcription factors. Identifying these transcription factors in crops will provide opportunities to tailor the senescence process to different environmental conditions and regulate the balance between yield and grain nutrient content. Here we use ten time points of gene expression data alongside gene network modelling to identify transcription factors regulating senescence in polyploid wheat. We observe two main phases of transcription changes during senescence: early downregulation of housekeeping and metabolic processes followed by upregulation of transport and hormone related genes. We have identified transcription factor families associated with these early and later waves of differential expression. Using gene regulatory network modelling alongside complementary publicly available datasets we identified candidate transcription factors for controlling senescence. We validated the function of one of these candidate transcription factors in senescence using wheat chemically-induced mutants. This study lays the ground work to understand the transcription factors which regulate senescence in polyploid wheat and exemplifies the integration of time-series data with publicly available expression atlases and networks to identify candidate regulatory genes.

## Introduction

Grain yield and nutrient content in cereal crops is determined by the accumulation of carbon, nitrogen and other nutrients in the grain towards the end of a plant’s life. The availability of these nutrients is strongly influenced by the process of senescence, a regulated developmental programme to remobilise nutrients from the vegetative tissues to the developing grain. Both the onset and rate of senescence influence grain yield and nutrient content. A delay in senescence may be associated with increased yield due to an extended period of photosynthesis (Gregersen et al., 2013; Thomas and Howarth, 2000). However, delayed senescence may also be associated with a decrease in grain nutrient content due to reduced nutrient remobilisation from green tissues (Distelfeld et al., 2014). Senescence is often associated with the visual loss of chlorophyll, however the initiation of senescence through signalling cascades, and early stages such as degradation of protein and RNA, are not visible (Buchanan-Wollaston et al., 2003; Fischer, 2012). Through these initial stages, and later during visual senescence, a programme of tightly-regulated changes occurs in gene expression (Buchanan-Wollaston et al., 2003; Fischer, 2012). Despite its importance, we know relatively little about the molecular control of senescence in crops such as wheat (Distelfeld et al., 2014).

This lack of knowledge is partly due to the difficulty of identifying genes regulating quantitative traits in the large wheat genome (IWGSC et al., 2018) as well as the subtle effects of individual gene copies (homoeologs) within the polyploid context (Borrill et al., 2015). These challenges mean that conventional genetic mapping approaches often take many years to identify causal genes. To date two genes have been identified to regulate senescence in wheat. The *NAM-B1* NAC transcription factor was identified to underlie a quantitative trait locus (QTL) for grain protein content and senescence (Uauy et al., 2006). A second NAC transcription factor, *NAC-S*, was found to have a strong correlation between its expression level and leaf nitrogen concentration in tandem with a role in regulating senescence (Zhao et al., 2015). However, to realise the potential to manipulate the rate and onset of senescence in wheat it will be necessary to gain a more comprehensive understanding of the network of transcription factors regulating this process. Identifying these transcription factors may enable the development of wheat varieties with a senescence profile tailored to maximise nutrient remobilisation whilst maintaining yield and providing adaption to local growing conditions.

The first step towards manipulating senescence at the molecular level is to understand the genes which are involved in the process, and the transcription factors which orchestrate gene expression changes during senescence. Over 50 % of micro and macronutrients remobilised to the developing grain originate from the uppermost (flag) leaf of the senescing wheat plant (Garnett and Graham, 2005; Kichey et al., 2007), making it a key tissue in which to understand the senescence process. Previous attempts have been made to characterise transcriptional changes in wheat flag leaves, however these studies have been either carried out with microarrays which were limited to a small set of 9,000 genes (Gregersen and Holm, 2007) or had a limited number of samples and time points (Pearce et al., 2014; Zhang et al., 2018). Decreases in the cost of RNA-Seq now mean that these constraints can be overcome through genome-wide expression studies across multiple time points. The recent publication of the wheat genome sequence with over 100,000 high confidence gene models (IWGSC et al., 2018) and accompanying functional annotations, enhances the ease and accuracy with which RNA-Seq data can be analysed in wheat. Systems biology approaches can start to make sense of the vast quantities of data produced and identify the regulatory pathways controlling quantitative traits (Kumar et al., 2015).

Our aim in this study was to identify the molecular pathways involved in senescence in wheat and determine candidate transcription factors controlling these processes in the flag leaf. We sequenced a ten time point expression timecourse of wheat senescence in the flag leaf from 3 days post anthesis until 26 days post anthesis which corresponded to the first signs of visual senescence. We identified the temporal progression of the senescence process at the molecular level and used gene regulatory network modelling to predict transcription factors which coordinate this developmental process. We confirmed the role of one of these candidate genes, *TraesCS2A02G201800* (*NAM-A2)*, in wheat itself.

## Results

### Growth and physiological measurements

To understand the transcriptional control of the initiation of senescence we harvested an early timecourse of senescence at 3, 7, 10, 13, 15, 17, 19, 21, 23 and 26 days after anthesis (DAA) (Figure 1A). SPAD chlorophyll meter readings in the flag leaf were maintained at a similar level from 3 to 21 DAA, with a significant decrease from 23 DAA (Figure S1). Percentage moisture of the grains decreased from 80.0 % at 3 DAA to 54.7 % at 26 DAA which corresponds to soft dough stage (Zadoks GS85 (Zadoks et al., 1974)) (Figure S2), indicating that the time period sampled included the majority of the grain filling period.

**Figure 1.**
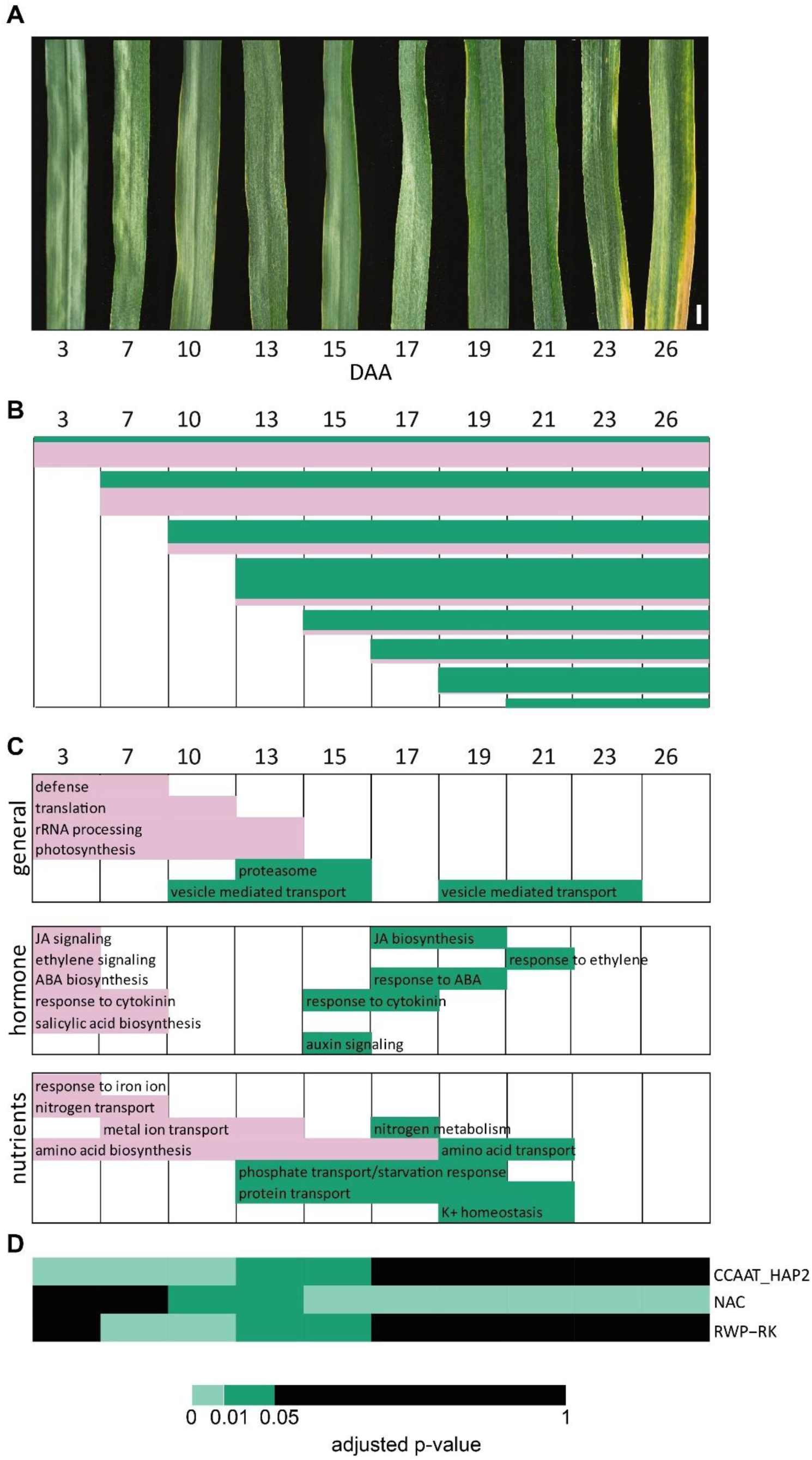
Transcriptional re-programming during flag leaf senescence. A) Timecourse of flag leaf senescence from 3 to 26 days after anthesis (DAA), scale bar represents 1 cm. B) Diagram showing representative patterns for genes which are consistently upregulated (green) or consistently downregulated (pink) during senescence (96.2 % of differentially expressed genes). Genes were grouped according to the first time of up or downregulation. The majority of genes in each pattern continued to be up or downregulated across the whole timecourse. Bar heights represent the number of genes in each expression pattern. The x axis represents time after anthesis, the axis is represented uniformly although time points are not evenly spaced. C) GO term enrichments are shown related to general, hormone and nutrient related processes. Filled rectangles represent that genes starting to be differentially expressed at that time point are enriched for that specific GO terms. Green rectangles represent upregulated genes, pink rectangles represent downregulated genes. D) Transcription factor families which were significantly enriched amongst upregulated genes across the timecourse. Significantly enriched time points are shown in green.

### Strong transcriptional changes occur during flag leaf senescence

RNA was extracted from the flag leaf blade with three replicates for each of the ten time points and sequenced. The RNA-Seq data was aligned to the RefSeqv1.1 transcriptome annotation (IWGSC et al., 2018) using kallisto (Bray et al., 2016). On average each sample had 38.7 M reads, of which 30.9 M mapped (78.9 %) (Table S1). We found that 52,905 high confidence genes were expressed at >0.5 transcripts per million (TPM) in at least one time point during flag leaf senescence, which corresponds to 49.0 % of high confidence genes. To identify genes differentially expressed during the timecourse, we used two programmes specifically designed for timecourse data: ImpulseDE2 (Fischer et al., 2018) and gradient tool (Breeze et al., 2011). In total 9,533 genes were identified as differentially expressed by both programmes, giving a high confidence set of differentially expressed genes. In addition, gradient tool identified at which time points the genes became differentially expressed which we used to determine the temporal changes in gene expression associated with senescence (Table S2).

To define the biological roles of these 9,533 genes we grouped them according to the first time point at which they were up or downregulated. For example, a gene first upregulated at 10 DAA was in group “U10” (up 10 DAA), whereas a gene first downregulated at this time point was assigned to group “D10” (down 10 DAA). Fewer than 4 % of genes were both up and down regulated during the timecourse and these were excluded from further analysis, resulting in 17 expression patterns (Table S2). In total approximately twice as many genes were upregulated during this senescence timecourse than downregulated (5,343 compared to 2,715). This indicates that senescence is actively regulated through transcriptional upregulation rather than a general downregulation of biological processes.

We found that the patterns of up and downregulation were not equally spaced throughout the timecourse. During the early stages of senescence the majority of differentially expressed genes were downregulated (825/1035 differentially expressed genes at 3 DAA), and these continued to be downregulated throughout the timecourse (Figure 1B). At the later stages of senescence relatively few genes started to be downregulated (e.g. 50 genes at 19 DAA). Instead the number of genes which started to be upregulated grew from 210 genes at 3 DAA to 1,324 genes at 13 DAA. After this peak of upregulation at 13 DAA, fewer genes started to be upregulated, although there were still over 500 genes upregulated at each of 15, 17 and 19 DAA. Genes which were upregulated even at early stages of senescence tended to continue to increase in expression level throughout the timecourse. At the latest stages of the timecourse when chlorophyll loss was visible, 23 and 26 DAA, very few genes started to be differentially expressed.

We found that this temporal divide into downregulation at the early stages of senescence and initiation of upregulation at the later stages of senescence was also reflected in different GO term enrichments in these groups of differentially expressed genes (Figure 1C; Table S3). The large numbers of genes which started to be downregulated at 3 and 7 DAA were enriched for GO terms relating to housekeeping functions (e.g. translation, photosynthesis and rRNA processing) as well as for central metabolic processes such as amino acid biosynthesis and starch biosynthesis. Alongside these housekeeping functions, downregulated genes were enriched for defence responses and hormone biosynthesis and signalling, indicating a reduction in the transcriptional responses to stimuli. Later in the timecourse, from 10 to 13 DAA, groups of genes started to be upregulated which were involved in vesicle mediated transport and the proteasome, indicating a remobilisation of components from the existing proteins. This is supported by the upregulation from 13 DAA of genes involved in phosphate and protein transport. From 15 DAA to 21 DAA waves of genes enriched for responses to cytokinin, ABA and ethylene were upregulated, indicating a temporal hierarchy of hormone responses during senescence.

To understand how these highly ordered and coordinated transcriptional changes are regulated we examined transcription factor (TF) expression patterns. We found that 2,210 TFs were expressed (> 0.5 TPM) during the timecourse but only 341 TFs (15.4 %) were differentially expressed. We calculated the percentage of differentially expressed TF per TF family across time (Figure 2). In general, each TF family tended to either be upregulated or downregulated as a whole (Figure 2), although there are exceptions such as the C2C2_CO-like and MADS_II family which showed upregulation and downregulation of different family members during the timecourse. Thus, the TFs which were downregulated during senescence largely belong to different TF families to those which were upregulated.

**Figure 2.**
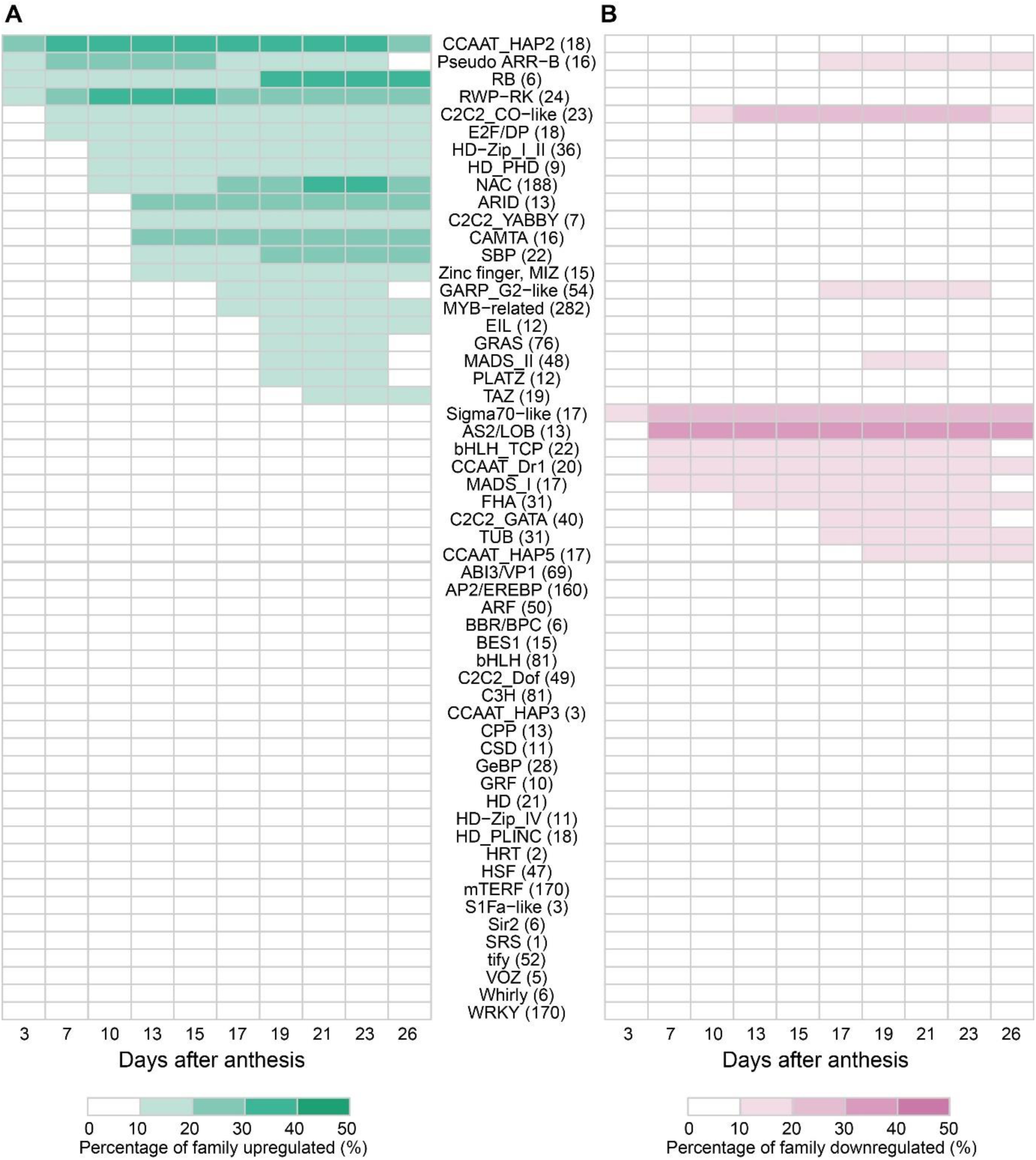
Percentage of expressed genes which were differentially expressed per transcription factor family at each time point. Upregulated (A) and downregulated (B) genes are shown. The total number of genes expressed in each family is shown in brackets after the family name.

While we observed a temporal gradient of TF families starting to be up and downregulated throughout the timecourse, we defined an initial (3 to 7 DAA) and later wave (13-19 DAA) when many TF families were up or downregulated. TF families which were upregulated in the initial wave from 3 to 7 DAA include the RWP-RK, pseudo ARR-B and CCAAT_HAP2 (NF-YA) families (Figure 2A). A distinct set of TF families were upregulated from 13 to 19 DAA in the later wave including CAMTA, GRAS and MADS_II. After these waves of upregulation were initiated, the same families tended to continue to be upregulated throughout the rest of the timecourse. Compared to all genes, the RWP-RK, CCAAT_HAP2 (NF-YA) and NAC families were significantly enriched (padj <0.01, Fisher test; Figure 1D) for upregulated genes at early (RWP-RK and CCAAT_HAP2 (NF-YA)) and late (NAC) time points. In all three families over 30 % of the expressed genes were upregulated during senescence corresponding to 61 NAC TFs (32.4 % of expressed NAC TFs) and eight RWP-RK and seven CCAAT_HAP2 (NF-YA) TFs (33.3 % and 38.9 % of expressed genes per family, respectively).

In parallel with certain TF families being upregulated, another group of TF families were downregulated during the senescence timecourse. The initial wave of downregulation largely occurred at 7 DAA and included the AS2/LOB, bHLH_TCP and MADS_I families. The later wave of downregulation initiated from 17 to 19 DAA and included the C2C2 GATA, GARP G2-like and MADS_II families. Similar to upregulation of TFs, the downregulation tended to continue throughout the rest of the timecourse, indicating a gradual change in transcription factor expression levels. None of the TF families were significantly enriched for downregulated genes compared to all genes.

These two waves of TF differential expression are analogous to the two waves of differential expression observed for all gene classes (Figure 1). This is consistent with TF roles as activators and repressors of gene expression. These results suggest that specific TF families initiate temporally distinct changes in gene expression, broadly classed into an initial (3 to 7 DAA) and later (13 to 19 DAA) response.

### Understand regulation using network modelling

Our results indicate that there are two main temporal waves of expression during senescence (3 to 7 DAA and from 13 to 19 DAA) which may be regulated by the associated upregulation of particular TF families. However, to understand the interactions between TFs and predict which ones may be key regulators (hub genes) driving this transcriptional programme we constructed a gene regulatory network. We used Causal Structure Inference (Penfold and Wild, 2011) which produces a directional network of transcription factor interactions. We included 213 TF which were both differentially expressed during the timecourse and which also had an expression level >5 TPM. We chose this threshold to maximise the number of informative genes, but to minimise noise by removing low expressed transcription factors which may not play a role in the transcriptional reprogramming of senescence.

To interpret the network it is necessary to determine the ‘edge weight threshold’ at which to include edges. Since our aim was to identify the most important TFs within the network to test as candidate genes for the regulation of senescence, we decided to compare the network across different thresholds. We hypothesised that by identifying TFs which were important across multiple thresholds we would be more likely to identify robust candidate genes. We found that from a threshold of 0.01 to 0.3 the number of edges reduced from 11,049 to 30 (Table 1). *NAM-A1*, a known regulator of senescence in wheat, was only present in the network at the lower thresholds of 0.01, 0.05 and 0.1. We therefore decided to focus on the networks which included *NAM-A1*, as it is likely that the more stringent thresholds (0.2 and 0.3) would also have excluded other TFs relevant to the senescence process. The other TF which had previously been identified to regulate senescence in wheat (*NAC-S*) was not detected as differentially expressed during our timecourse so it was not used to construct the network or determine appropriate thresholds.

**Table 1.** Comparing CSI network at different thresholds for edge weight.

| Edge weight threshold | # genes | # edges | *NAM-A1* |
| --- | --- | --- | --- |
| 0.01 | 213 | 11,049 | Yes |
| 0.05 | 204 | 692 | Yes |
| 0.1 | 132 | 182 | Yes |
| 0.2 | 67 | 59 | No |
| 0.3 | 38 | 30 | No |

We determined the importance of a gene within the network using two measures: ‘edge count’ which is the number of connections to other genes, and ‘betweenness centrality’ which is a measure of the number of shortest paths which pass through that gene and represents a measure of how essential the gene is to the flow of information around the network. We calculated percentage rankings for genes in each the three thresholds (0.01, 0.05 and 0.1) according to their edge count and betweenness centrality to allow comparison across networks with different numbers of genes. We found that 24.7 % of genes (53 genes) were ranked in the top 20 % of genes in at least one threshold for betweenness centrality and one threshold for edge count (Figure 3A). We consider these to represent good candidate genes for further investigation. Amongst the 53 top ranked genes we found that three transcription factor families were enriched compared to all 213 transcription factors in the network: GARP_G2-like, HSF and RWP-RK (χ^2^ < 0.01, 0.05 and 0.001 respectively; Figure 3B-D). Interestingly the RWP-RK family was also significantly enriched for upregulation during senescence (Figure 1D), in addition to being enriched amongst top ranked genes in the network. One family was significantly depleted in the top ranked genes: the WRKY family (χ^2^ < 0.05) which was surprising since WRKY transcription factors have been reported to regulate senescence in rice (Muho et al., 2014) and Arabidopsis (Phukan et al., 2016).

**Figure 3.**
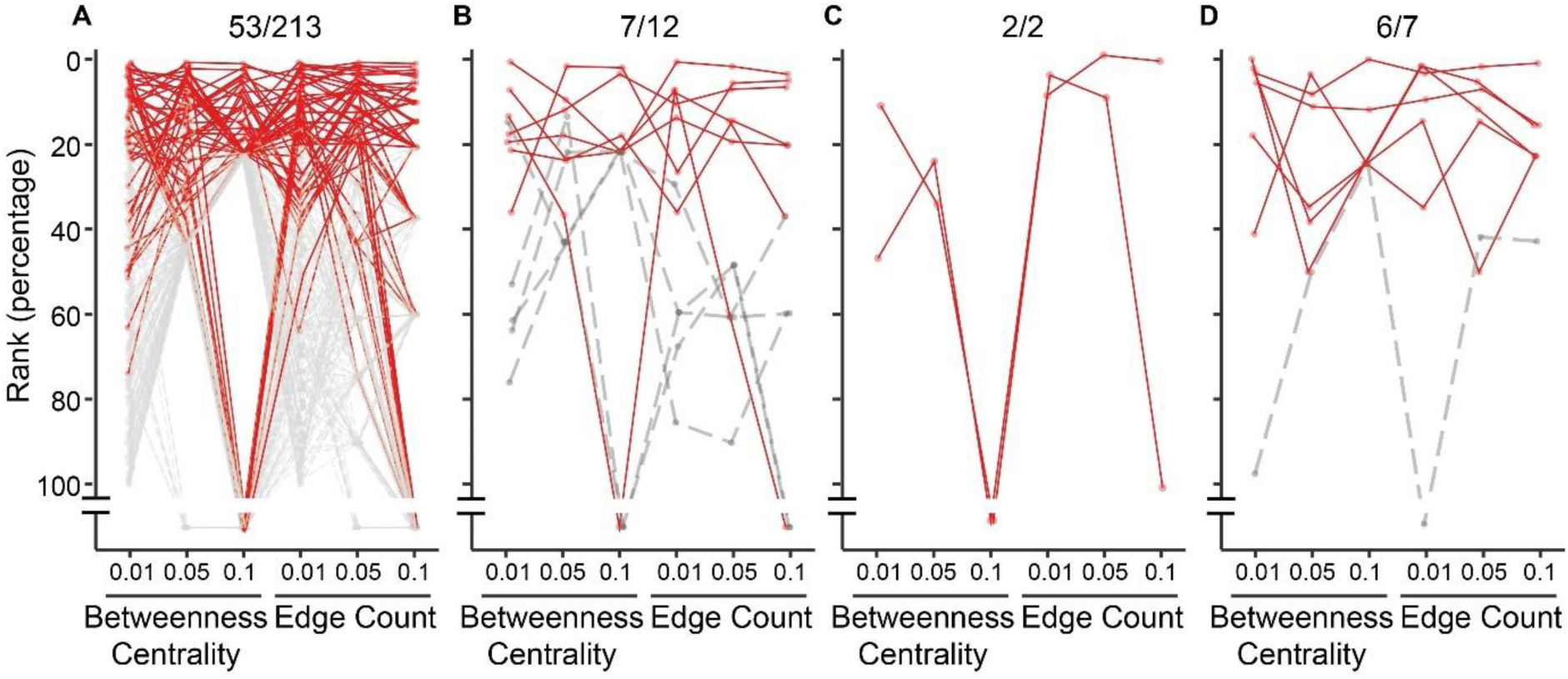
Comparisons of percentage rankings for betweenness centrality and edge count. A) All 213 genes in the network are shown with their percentage ranking at each threshold. The 53 genes ranked in the top 20 % across at least one betweenness centrality and one edge count threshold are shown in red. B) GARP_G2-like, C) HSF and D) RWP-RK TF families enriched within the top ranked 53 genes. B-D) Genes within the top 53 genes are shown in red, whilst genes in these families not in the top 53 genes are shown in grey (there are no HSF outside the top 53 genes). Numbers above graphs indicate the number of TFs in the top 20 % out of the total number of TFs shown in the graph. Genes not present in the network at higher thresholds are represented as points below the Y-axis break.

### Independent data supporting candidate gene prioritisation

Although we could prioritise candidates based on information solely from the network, we decided to also incorporate other data sources to help distinguish the 53 top ranked candidate genes. We focused on three additional datasets: 1) expression data from an independent experiment with 70 tissues/time points in the spring wheat variety Azhurnaya which included senescing leaves (Ramirez-Gonzalez et al., 2018), 2) a GENIE3 network of predicted transcription factors – target relationships from 850 independent expression samples (Ramirez-Gonzalez et al., 2018) and 3) information from orthologs in Arabidopsis and rice.

We found that 16 out of the 53 top ranked genes from the network were expressed over two-fold higher in senescing tissues than in other tissues across the Azhurnaya developmental experiment (Figure 4A). This independent dataset suggests that these 16 genes may play a specific role in senescing tissues and we hypothesise that they would be less likely to induce pleiotropic effects when their expression is altered in mutant or transgenic lines. We also tested whether these 53 candidate genes had targets which were predicted to play a role in senescence, using the independent GENIE3 transcription factor - target network. We found that five of the candidate genes had targets enriched for senescence-related GO terms (Figure 4B), however this did not include *NAM-A1*, which suggested this approach might miss some interesting candidate genes. Therefore, we also tested whether the candidate genes had any shared targets genes with *NAM-A1* which might indicate they act together in the same senescence related pathway. We found that 22 genes had one or more shared target genes with *NAM-A1* (Figure 4B). Even an overlap of one target gene is significantly more than the expected zero overlap between *NAM-A1* and a random transcription factor (Sign test, p-value < 0.001). We found that only two of the candidate genes were direct targets of *NAM-A1*, and these included *NAM-D1*, the D-genome homoeolog of *NAM-A1*, and *NAM-A2*, an uncharacterised paralog of *NAM-A1* which is located on a different chromosome.

**Figure 4.**
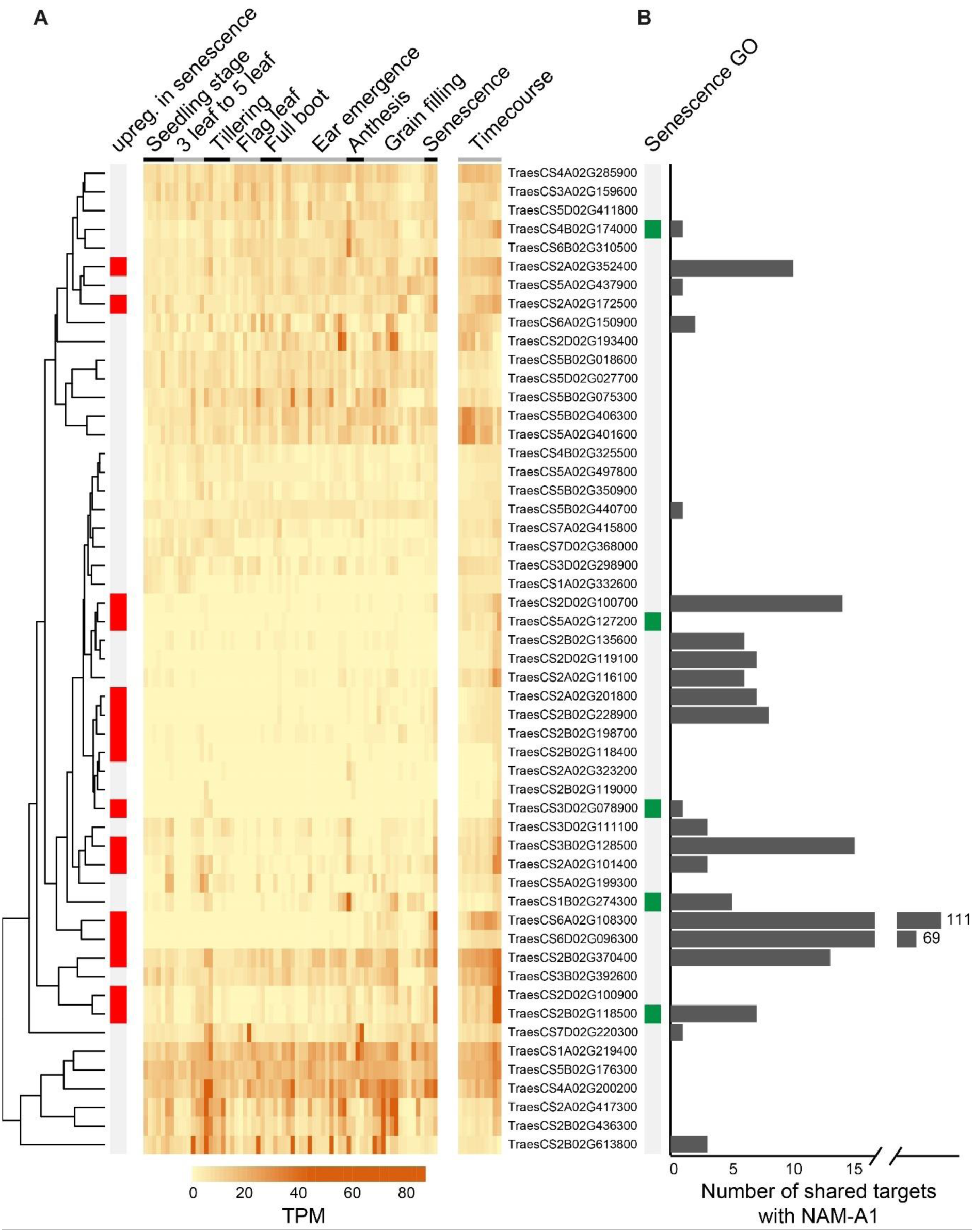
Additional information for 53 top ranked candidate genes. A) Expression from an independent RNA-Seq experiment using Azhurnaya spring wheat (left part of heatmap) and expression in the senescence timecourse (right part of heatmap, “Timecourse”). Each of the 53 genes is represented in one row, and rows are sorted according to the similarity of the expression patterns (dendrogram to left). Genes which were over two-fold upregulated in senescence compared to other tissues/time points in Azhurnaya are highlighted by red boxes (“upreg. in senescence”). Expression level is measured in transcripts per million (TPM). B) Targets of the TF predicted by independent GENIE3 network. Genes with downstream targets enriched for senescence GO terms in the independent GENIE3 network are marked with green boxes. The bar graph shows the number of shared targets with *NAM-A1*. *NAM-A1* (*TraesCS6A02G108300*) has 111 targets and its homoeolog *NAM-D1* (*TraesCS6D02G096300*) has 69 shared targets, shown with broken axis.

We identified Arabidopsis and rice orthologs for the 53 candidate genes using *Ensembl*Plants gene trees (Kersey et al., 2018). Arabidopsis orthologs were identified for 45 genes, and rice orthologs for 46 genes (Table S4). The Arabidopsis orthologs of thirteen genes had known leaf senescence functions, these corresponded to seven Arabidopsis genes in total due to several wheat homoeologs sharing the same ortholog. The rice orthologs of these genes had not been reported to have a leaf senescence function because the majority had not been phenotypically characterised. In addition, a large proportion of both Arabidopsis and rice orthologs (orthologs of thirteen and nine wheat genes respectively) play roles in nitrogen responses, consistent with the tight coordination expected between senescence and nitrogen remobilisation from flag leaves.

### Validation of candidate gene *NAM-A2*

Using the additional information sources above we selected *NAM-A2* (*TraesCS2A02G201800*) for phenotypic characterisation in wheat because it was amongst our 53 top ranked candidate genes, was upregulated in senescing leaves and shared many downstream target genes with *NAM-A1*. Furthermore the *NAM-A2* homoeolog, *NAM-B2* (*TraesCS2B02G228900*) was also amongst the top 53 candidate genes. *NAM-A2* is a closely related paralog of *NAM-A1* which regulates senescence and nutrient remobilisation (Avni et al., 2014; Uauy et al., 2006). The homoeolog of *NAM-A2*, *NAM-B2*, was previously found to cause a slight delay in senescence (Pearce et al., 2014) but *NAM-A2* has not been previously characterised so was a strong candidate as a transcription factor which might regulate senescence.

To test the predictions of our model we identified TILLING mutations in *NAM-A2* and *NAM-B2* in a tetraploid Kronos background (Krasileva et al., 2017; Uauy et al., 2009). Due to the potential redundancy between homoeologs in wheat (Borrill et al., 2015) we decided to generate double *NAM-A2*/*NAM-B2* mutants through crossing. We identified a mutation leading to a premature stop codon in *NAM-B2* (R170*; between subdomains D and E of the NAC domain (Kikuchi et al., 2000)), which is predicted to abolish protein function by creating a truncated protein lacking part of the NAC DNA binding domain. For *NAM-A2* we could not identify any mutations which would cause truncations, instead we selected three missense mutations which were in highly conserved domains and were thus expected to play important roles in protein function (Figure 5A). These were located in the A, C and D NAC subdomain and were predicted to be highly deleterious according to SIFT and PSSM scores. We crossed each of the *NAM-A2* missense mutants to the *NAM-B2* truncation mutant to create segregating populations from which wild type, single and double mutants which were phenotyped in the F3 generation.

**Figure 5.**
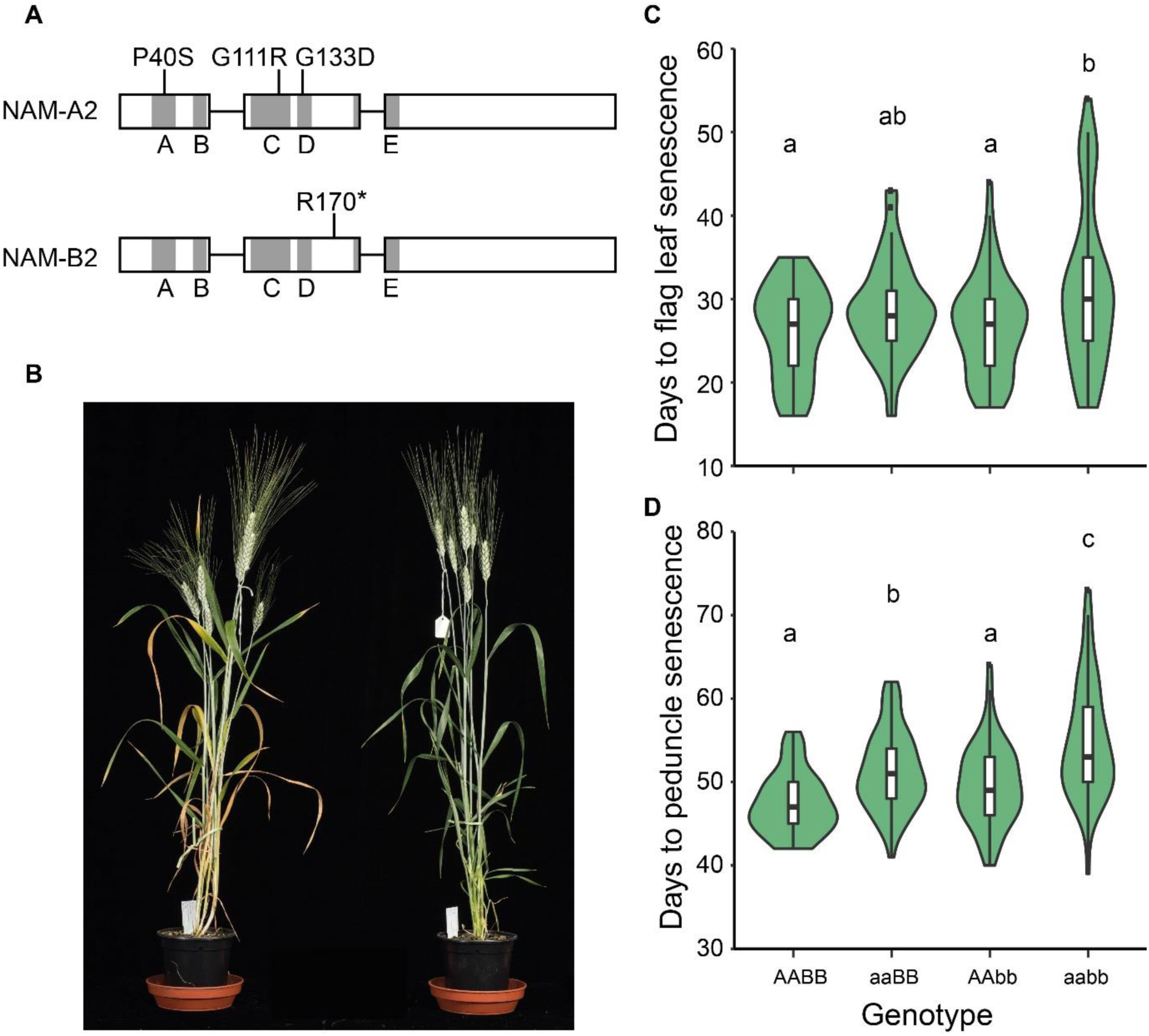
Mutants in *NAM-A2* and *NAM-B2*. A) Selected missense mutations in *NAM-A2* and stop mutation in *NAM-B2*. Grey regions are the NAC subdomains A-E. Subdomain E spans the end of exon 2 and the start of exon 3. B) Wild type sister line (left) and *NAM-A2 NAM-B2* double homozygous (aabb) mutant (right), 37 days after anthesis. C) Days from heading to flag leaf senescence and D) days from heading to peduncle senescence in wild type, single and double mutants. Letters indicate significant differences p < 0.05, with ANOVA post-hoc Tukey HSD.

Across the three populations with different missense mutations in *NAM-A2*, and a common truncation mutation in *NAM-B2*, there was a significant delay of 4.9 days in flag leaf senescence in the double mutant compared to wild type (padj <0.01, ANOVA post-hoc Tukey HSD; Figure 5B-C). There were no significant differences between the single mutants and wild type in flag leaf senescence. Peduncle senescence was significantly delayed by 7.4 days in the double mutant compared to wild type (padj <0.001, ANOVA post-hoc Tukey HSD; Figure 5D), and in addition the single A mutant was significantly later in peduncle senescence than wild type (3.9 days, padj <0.001, ANOVA post-hoc Tukey HSD). The single B mutant was not significantly different from wild type suggesting that the A genome homoeolog has a stronger effect on senescence than the B genome homoeolog. Since the comparison is between different types of mutations (missense compared to a truncation mutation) interpretation of the relative magnitudes is difficult, although the truncation mutation in the B genome would have been expected to produce at least an equivalent effect to the missense mutation in the A genome. These effects were largely consistent across the three different missense mutations, although the mutation in subdomain C (G111R) had the largest effect when combined into a double mutant compared to wild type (Figure S3).

## Discussion

In this work we have characterised the transcriptional processes associated with senescence in the wheat flag leaf. We found that specific transcription factor families are associated with these changes in transcription and have used gene regulatory network modelling, alongside additional complementary information, to identify candidate genes controlling this process. We confirmed that one of these genes, *NAM-A2*, plays a role in senescence in wheat itself.

### Time-resolved transcriptional control of senescence in wheat

We found that although 52,905 genes were expressed in senescing flag leaves, only 9,533 genes were differentially expressed during this time period. Sampling ten time points allowed us to observe that these 9,533 differentially expressed genes were largely divided into two temporal waves of transcriptional changes which may not have been captured using a less time-resolved set of data. Frequent sampling has also proved informative in other time dependent processes in wheat such as pathogen infection (Dobon et al., 2016) and represents a powerful approach to understand the coordination and regulation of gene expression changes throughout development and environmental responses (Bar-Joseph et al., 2012; Lavarenne et al., 2018).

We found that during the first wave of transcriptional changes the majority of differentially expressed genes were downregulated, and these groups were enriched for GO terms related to translation, photosynthesis and amino acid biosynthesis. During the second wave, genes started to be upregulated with enrichment for GO terms related to vesicle mediated transport, protein transport and phosphate transport. The chronology of biological processes is well conserved with Arabidopsis. For example early downregulation of chlorophyll related genes is observed in both Arabidopsis (Breeze et al., 2011) and wheat, whilst transport processes are upregulated later during senescence. The temporal order of senescence related processes is also broadly conserved in maize (Zhang et al., 2014) and rice (Lee et al., 2017).

The importance of transcription factors to tightly coordinate the transcriptional changes happening during the senescence is well known from other plant species (Podzimska-Sroka et al., 2015; Woo et al., 2016). We found that particular TF families were up and downregulated in two distinct waves, an initial and later response, following the pattern for all differentially expressed genes. We found that three transcription factor families were enriched for upregulated genes during senescence at early (CCAAT_HAP2 and RWP-RK) and late (NAC) stages. Members of the NAC family have been characterised to play a role in regulating senescence in both wheat (Uauy et al., 2006; Zhao et al., 2015) and other plant species (Podzimska-Sroka et al., 2015). The CCAAT_HAP2 (NF-YA) family is less well characterised in this process but one member has been shown to delay nitrate-induced senescence in Arabidopsis (Leyva-González et al., 2012). The RWP-RK family is known in Arabidopsis to control nitrogen responses (Chardin et al., 2014), and in cereals nitrogen remobilisation is closely connected with senescence highlighting the potential for further investigations into this family in the future. Surprisingly the WRKY transcription factor family, which has been reported to play important roles in senescence in several other species such as Arabidopsis (Breeze et al., 2011; Woo et al., 2013), cotton (Lin et al., 2015) and soybean (Brown and Hudson, 2017), was not enriched for upregulation during senescence in wheat. It is possible that relatively few members of the WRKY family function in regulating senescence in wheat or that the function of WRKY TFs has diverged between wheat and other plant species. This potential for divergence in the regulation of senescence between species is supported by experiments characterising the rice ortholog of *NAM-B1*. Whilst the *NAM-B1* transcription factor in wheat regulates monocarpic senescence, the ortholog in rice (*Os07g37920*) regulates anther dehiscence and does not affect monocarpic senescence (Distelfeld et al., 2012).

### Identifying candidate genes in networks

One of the aims of this study was to identify transcription factors which regulate the process of senescence. The rationale behind this approach was that transcription factors control other genes and therefore may have a strong and readily detectable effect on the process of senescence. Secondly, in crops, transcription factors have been frequently selected under QTLs for important traits such as flowering time (*PPD1*, *VRN1*) (Beales et al., 2007; Yan et al., 2003) and cold tolerance (*CBF*) (Knox et al., 2008) due to their strong phenotypic effects. Thus, identified candidate transcription factors regulating senescence might also prove to be useful breeding targets.

Through examining the expression patterns of transcription factors in detail we identified transcription factor families which were enriched for upregulation during senescence, however this analysis cannot provide information about which of the individual transcription factors within the family might be more important in regulating the senescence process. To address this question, we used Causal Structure Inference (Penfold and Wild, 2011) to identify interactions between transcription factors. Our hypothesis was that central transcriptional regulators of senescence would regulate other transcription factors to create a regulatory cascade to influence the thousands of genes differentially expressed during senescence. Amongst the 53 top ranked transcription factors in the network, three TF families were enriched: the GARP-G2-like, HSF and RWP-RK. Members of the GARP_G2-like family have been reported to play a role in senescence in rice (Rauf et al., 2013). HSF transcription factors are associated with stress responses, and although no members have been associated with developmental senescence, stress responsive genes are also closely associated with environmentally-induced senescence, and common regulation has been observed in Arabidopsis (Woo et al., 2013). The RWP-RK family is of interest because it also significantly enriched for upregulation during senescence, in addition to being enriched amongst top ranked genes in the network. This adds further weight to the hypothesis that the RWP-RK TFs may play a role in senescence, in addition to their known role in nitrogen responses. The roles of these identified TF can now be directly tested in wheat to determine whether they regulate senescence using gene editing and TILLING (Borrill et al., 2015).

To further delimit this list of candidate genes we used information from independent datasets (developmental timecourse of expression and GENIE3 TF-target network) to prioritise candidate genes. The approach to combine additional data sets was also applied in Arabidopsis where a Y1H screen was used in conjunction with Causal Structure Inference to help to identify regulatory interactions in senescence and pathogen infection (Hickman et al., 2013). Another approach which can be used to narrow down candidate genes is to examine how the network is perturbed in transcription factor mutants. This approach was used in Arabidopsis to identify three NAC transcription factors which regulate senescence (Kim et al., 2018) and could now be applied in wheat using the TILLING mutant resource (Krasileva et al., 2017), for example starting with the mutants generated in this study.

To test the predicted function of these candidate genes in regulating wheat senescence, we focused on *NAM-A2*, which is a paralog of the known *NAM-B1* gene. We found significant delays in flag leaf and peduncle senescence in *NAM-A2/NAM-B2* double mutants, indicating that the genes predicted by the network play roles in senescence. The peduncle senescence phenotype indicates that this approach can identify genes which regulate senescence across different tissues, not only in the flag leaf, and may reflect that monocarpic senescence in wheat is a developmental process regulated across the whole plant. Ongoing work is currently characterising the additional candidate genes through the development of wheat double mutants for phenotypic characterisation.

### Future directions

This study has uncovered candidate transcription factors which may regulate senescence in wheat and has confirmed the role of one of these genes in regulating senescence. It will be of great interest to determine whether these genes control only senescence or also affect nutrient remobilisation and hence influence final grain nutrient content. In addition to deepening our understanding of the molecular regulation of senescence, this study lays the ground work to use this network-enabled approach to identify transcription factors regulating a range of different biological processes which happen across a timecourse. This approach is not only applicable to developmental processes but could equally be applied to abiotic and biotic stresses, as has been carried out in other plant species (Hickman et al., 2013). This approach could also be applied to identify candidate genes for traits in species without genome sequences, although a transcriptome would need to be assembled from the RNA-Seq data. The advent of genome-editing means that the prediction of gene function could readily be tested in any transformable species.

## Conclusion

The availability of a fully sequenced reference genome for wheat, alongside functional genomic resources such as the TILLING population, have brought wheat biology into the genomics era and have made possible studies which even a few years ago would have been unthinkable. Here we have used these new resources to characterise the transcriptional processes occurring during wheat senescence. We found that specific transcription factor families are associated with this process in wheat, some of which have been reported in other species, but others present new links between transcription factor families and the process of senescence. Although these associations do not prove causality, the hypotheses generated can now be tested experimentally in wheat using TILLING or gene editing. Gene network modelling, when used in conjunction with complementary datasets, is a powerful approach which can accelerate the discovery of genes regulating biological processes in both model and crop species.

## Methods

### Plant growth for RNA-Seq timecourse

We pre-germinated seeds of hexaploid wheat cv. Bobwhite on moist filter paper for 48 h in 4 °C followed by 48 h in the dark at room temperature. These pre-germinated seeds were sown in P40 trays in 85% fine peat with 15% horticultural grit. Plants were potted on at 2–3 leaf stage to 1L square pots with 1 plant per pot in Petersfield Cereal Mix (Petersfield, Leicester, UK). Plants were grown in 16 h light at 20 °C, with 8 h dark at 15 °C. The main tiller was tagged at anthesis, and the anthesis date was recorded.

#### Phenotyping for RNA-Seq timecourse

We measured the chlorophyll content of flag leaves across the timecourse from 3 to 26 days after anthesis (DAA) using a SPAD-502 chlorophyll meter (Konica Minolta). The time points used were 3, 7, 10, 13, 15, 17, 19, 21, 23 and 26 DAA. We measured the flag leaf from the main tiller (tagged at anthesis) for five separate plants for each time point, taking measurements at 8 different locations distributed along the length of each flag leaf. Three of these measured leaves were subsequently harvested for RNA extraction.

We measured the grain moisture content across the timecourse from 3 to 26 days after anthesis, using the same time points as for chlorophyll measurements. We harvested eight grains from central spikelets (floret positions 1 and 2) within the primary spike of five separate plants at each time point, these grains were weighed, and then dried at 65 °C for 72 hours before re-weighing. The difference in weight was used to calculate the percentage grain moisture content.

### Tissue harvest, RNA extraction and sequencing

#### Harvesting

The flag leaf from the main tiller was harvested at 3, 7, 10, 13, 15, 17, 19, 21, 23 and 26 DAA from three separate plants (three biological replicates). We harvested the middle 3 cm of the flag leaf lengthways to have a region of the leaf which was synchronised in its developmental stage. We flash froze the samples in liquid nitrogen, then stored them at -80 °C prior to processing. In total we harvested 30 samples.

#### RNA extraction

We ground the samples to a fine powder in mortar and pestles which had been pre-chilled with liquid nitrogen. We extracted RNA using Trizol (ThermoFisher) according to the manufacturer’s instructions, using 100 mg ground flag leaf per 1 ml Trizol. We removed genomic DNA contamination using DNAseI (Qiagen) according to the manufacturer’s instructions and cleaned up the samples using the RNeasy Mini Kit (Qiagen) according to the manufacturer’s instructions.

#### Library preparation

The quality of the RNA was checked using using a Tecan plate reader with the Quant-iT™ RNA Assay Kit (Life technologies/Invitrogen Q-33140) and also the Quant-iT™ DNA Assay Kit, high sensitivity (Life technologies/Invitrogen Q-33120) Finally the quality of the RNA was established using the PerkinElmer GX with a high sensitivity chip and High Sensitivity DNA reagents (PerkinElmer 5067-4626). Thirty Illumina TruSeq RNA libraries were constructed on the PerkinElmer Sciclone using the TruSeq RNA protocol v2 (Illumina 15026495 Rev.F). After adaptor ligation, the libraries were size selected using Beckman Coulter XP beads (Beckman Coulter A63880). This removed the majority of un-ligated adapters, as well as any adapters that may have ligated to one another. The PCR was performed with a PCR primer cocktail that annealed to the ends of the adapter to enrich DNA fragments that had adaptor molecules on both ends. The insert size of the libraries was verified by running an aliquot of the DNA library on a PerkinElmer GX using the High Sensitivity DNA chip and reagents (PerkinElmer CLS760672) and the concentration was determined by using the Tecan plate reader.

#### Sequencing

The TruSeq RNA libraries were normalised and equimolar pooled into one final pool using elution buffer (Qiagen). The library pool was diluted to 2 nM with NaOH and 5μL transferred into 995μL HT1 (Illumina) to give a final concentration of 10pM. 120 μL of the diluted library pool was then transferred into a 200 μL strip tube, spiked with 1% PhiX Control v3 and placed on ice before loading onto the Illumina cBot. The flow cell was clustered using HiSeq PE Cluster Kit v3, utilising the Illumina PE_Amp_Lin_Block_Hyb_V8.0 method on the Illumina cBot. Following the clustering procedure, the flow cell was loaded onto the Illumina HiSeq 2000/2500 instrument following the manufacturer’s instructions. The sequencing chemistry used was HiSeq SBS Kit v3 with HiSeq Control Software 2.2.58 and RTA 1.18.64. Reads (100 bp, paired end) in bcl format were demultiplexed based on the 6bp Illumina index by CASAVA 1.8, allowing for a one base-pair mismatch per library, and converted to FASTQ format by bcl2fastq.

### RNA-Seq data analysis

#### Mapping

We pseudoaligned the samples using kallisto v0.44.0 with default parameters to the RefSeqv1.0 annotation v1.1 (IWGSC et al., 2018). Transcripts per million (TPM) and counts for all samples were merged into a single dataframe using tximport v1.0.3 (Soneson et al., 2016). Scripts for data analysis are available from https://github.com/Borrill-Lab/WheatFlagLeafSenescence.

#### Differential expression analysis

We filtered for high confidence genes which were expressed on average >0.5 TPM in at least one time point; this excluded low expressed genes and low confidence gene models from further analysis, consistent with previous analyses in wheat (Ramirez-Gonzalez et al., 2018). In total 52,905 genes met this condition. We used the count expression level of these genes for differential expression analysis using the R package ImpulseDE2 v1.4.0 (Fischer et al., 2018), all counts were rounded to the nearest integer before they were analysed with ImpulseDE2. In parallel we used the TPM expression level of these 52,905 genes for differential expression analysis using Gradient Tool v1.0 (Breeze et al., 2011) with the normalisation enabled on Cyverse (https://de.cyverse.org/de/) (Merchant et al., 2016). To select a high confidence set of differentially expressed genes we only retained genes which were differentially expressed padj <0.001 from ImpulseDE2 and which were differentially expressed according to Gradient Tool with a z-score of >|2|. We grouped the 9,533 high confidence differentially expressed genes according to the first time point at which they were up or downregulated. For example, a gene first upregulated at 10 DAA was in group “U10” (up 10 DAA), whereas a gene first downregulated at this time point was assigned to group “D10” (down 10 DAA). Genes which were both up and downregulated during the timecourse (<4 % of all differentially expressed genes) were grouped according to the time point of first differential expression with the opposite change also indicated. For example a gene upregulated at 10 DAA and then downregulated at 15 DAA was grouped as U10D (the second time point of differential expression was not recorded in the grouping). These groupings are available in (Table S2). The minority of genes with both up and downregulation (<4 %of all differentially expressed genes) were excluded from further analysis.

#### GO term enrichment

We obtained GO terms from the RefSeqv1.0 annotation and transferred them from the annotation v1.0 to v1.1. We only transferred GO terms for genes which shared >99 % identity across > 90% of the sequence (105,182 genes; 97.5 % of all HC genes annotated in v1.1). GO term enrichment was carried out for each group of differentially expressed genes (groups defined according to the first time point at which genes were upregulated or downregulated, see above) using GOseq v1.24.0.

#### TF annotation

Genes which were annotated as TFs were obtained from https://opendata.earlham.ac.uk/wheat/under_license/toronto/Ramirez-Gonzalez_etal_2018-06025-Transcriptome-Landscape/data/data_tables/ (Ramirez-Gonzalez et al., 2018).

#### Gene regulatory network construction

We selected transcription factors which were amongst the 9,533 differentially expressed genes. We filtered to only keep transcription factors which were expressed on average >5 TPM in at least one time point. We used the TPM gene expression values as input to Causal Structure Inference (CSI) v1.0 (Penfold and Wild, 2011) which was run through Cyverse (https://de.cyverse.org/de/) (Merchant et al., 2016). The parameters used with CSI were the defaults (parental set depth =2, gaussian process prior = 10;0.1, weight truncation = 1.0E-5, data normalisation = standardise (zero mean, unit variance), weight sampling = FALSE). The output marginal file was converted to Cytoscape format using hCSI_MarginalThreshold v1.0 in Cyverse with a probability threshold of 0.01. We used this file for directed network analysis in Cytoscape v3.6.1 (Shannon et al., 2003) which produced network statistics. We used Cytoscape to filter the network for edge count and betweenness centrality at 0.01, 0.05, and 0.1.

#### GENIE3 data

We identified the targets of TF using a TF-target network which was previously published (Ramirez-Gonzalez et al., 2018). Only connections amongst the top one million links were considered in this analysis. The network had been produced by a random forest approach (GENIE3) (Huynh-Thu et al., 2010) using 850 RNA-Seq samples.

#### Ortholog identification

We identified the rice and Arabidopsis orthologs of the wheat genes using *Ensembl*Plants gene trees (Kersey et al., 2018). In cases where relationships were not one to one, all possible paralogous copies were included in the analysis.

#### Visualisation

Graphs were made in R using the packages ggplot2 (Wickham, 2016), NMF (aheatmap function) (Gaujoux and Seoighe, 2010) and pheatmap (Kolde, 2013).

### Candidate gene validation

#### Phenotyping of NAM-2 mutants

We selected mutant lines from the Kronos TILLING population (K0282, K0427, K3240) (Krasileva et al., 2017) with missense mutations (G111R, G133D, P40S, respectively) in *NAM-A2* (*TraesCS2A02G201800*). These *NAM-A2* mutant lines were crossed with a line containing a mutation inducing a premature stop codon in *NAM-B2* (*TraesCS2B02G228900*) (K4452; R170*). For each of the three crosses, heterozygous F1 seeds (AaBb) were self-pollinated to produce an F2 population. We selected double homozygous mutant (aabb), single homozygous mutant (aaBB or AAbb) and double homozygous wild type plants (AABB) in the F2. Seeds from two individuals of each genotype in the F2 population were grown in greenhouse conditions for phenotyping from Jan 2018 – May 2018 in Norwich with 16 h supplemental lighting and a daytime temperature of 18 °C, and a night-time temperature of 12 °C. We tagged the main tiller at anthesis and recorded the anthesis date. We scored flag leaf senescence as the date when the flag leaf of the main tiller had lost chlorophyll from 25 % of the flag leaf blade. We scored peduncle senescence as the date when the top 3 cm of the peduncle lost all green colour and turned straw-yellow.

## Data availability

RNA-Seq raw reads have been deposited in the SRA accession PRJNA497810. Scripts for data analysis are available from https://github.com/Borrill-Lab/WheatFlagLeafSenescence.

## Additional files

Figures S1 – S3.

Tables S1-S4.

## Author contributions

PB and CU conceived, designed and coordinated the study. PB harvested tissue for the timecourse and collected the associated chlorophyll and grain moisture content phenotypic data. PB carried out the RNA extraction, analysed the RNA-Seq data and built the gene regulatory network model. PB identified mutations in *NAM-A2* and *NAM-B2* for crossing and designed KASP markers. JS carried out crossing of *NAM-A2* and *NAM-B2* mutant lines. JS and PB carried out KASP genotyping. PB and SH carried out phenotyping of the *NAM2* mutant lines. PB wrote the manuscript. CU, SH and JS edited the manuscript.

## Acknowledgments

This work was supported by the UK Biotechnology and Biological Sciences Research Council (BBSRC) through an Anniversary Future Leader Fellowship to P.B. (BB/M014045/1) and the Designing Future Wheat (BB/P016855/1) and GEN (BB/P013511/1) ISPs. S.A.H. was supported by the John Innes Foundation. This research was also supported in part by the NBI Computing infrastructure for Science (CiS) group through the HPC resources.

